# Genetically induced mouse model for colon-specific epithelial cell tumorigenesis driven by loss of K8 and Apc

**DOI:** 10.1101/2025.02.03.635623

**Authors:** Mina Tayyab, Mira M. E. Minkkinen, Carl-Gustaf A. Stenvall, Lauri Polari, Yatrik M. Shah, Diana M. Toivola

## Abstract

Loss of keratin 8 (K8) has been shown to increase susceptibility towards colonocyte hyperproliferation and tumorigenesis. However, most colorectal cancer (CRC) mouse models require carcinogen, develop small intestinal tumors or have long latency period. The aim was to establish a genetic, colon-specific and more human like CRC model driven by loss of K8 and Apc. Colon epithelium specific targeting using the CDX2P-CreER^T2^ mice was used to generate K8^flox/flox^; CDX2P-CreER^T2^ and K8^flox/flox^; CDX2P-CreER^T2^; Apc^flox/+^ mice. Body weight and stool consistency were monitored, and colon was analyzed for tumor burden and histopathology. Keratin expression, inflammation, and proliferation were assessed using immunoblotting and immunofluorescence analysis. This data was compared to K8 expression analysis in patients with CRC using UALCAN database. K8 downregulation in adult K8^flox/flox^; CDX2P-CreER^T2^ mice triggers mild diarrhea and leads to loss of K8 and reduced partner keratin levels in a mosaic pattern in the colonic epithelium, while ileal K8 protein levels are unchanged. K8-negative colon areas display increased crypt loss and more MPO+ cells predominantly in the proximal colon. Increased colonocyte proliferation is observed as increased percentage of Ki67+ cells and lower IL-22BP protein levels throughout the colon. These mice with additional monoallelic Apc inactivation show increased colon tumor formation. In colon adenocarcinoma patients, K8 expression is decreased independent of disease type and stage, age or gender. New genetic and colon-specific mouse model with loss of K8 and Apc adequately resembles human CRC. This study also highlights a role of colonocyte K8 in maintaining colon epithelial integrity and protecting against colon tumorigenesis.

## Introduction

The intestinal epithelium serves as a selectively permeable barrier and exhibits the highest proliferation rate of any tissue, with complete self-renewal occurring approximately every 3–5 days in mammals ^1,2^. Dysregulation of this tightly controlled barrier and rapid cell turnover often predisposes individuals to life-threatening conditions, including inflammatory bowel disease (IBD) ^3^ and colorectal cancer (CRC) ^4^. CRC is a multifactorial malignancy, and its incidence is rising rapidly, particularly among young adults ^5^. Early detection of CRC and identification of the factors driving its pathogenesis are crucial for guiding treatment strategies. In this regard, experimental models have significantly advanced our understanding of CRC and served as a bridge between preclinical and clinical research. Mouse models are extensively used to study CRC pathogenesis and explore treatment options ^6^. However, these models have several limitations. Many require carcinogenic chemicals, such as azoxymethane (AOM), and/or take a long time to develop tumors ^7^. Additionally, the small intestinal tumor location in Apc inactivation-based models is notably different from that in humans, where the primary tumor site is typically the colorectum ^8^. Therefore, there is a need to develop models that more accurately recapitulate human CRC.

In efforts to refine Apc-based mouse models, use of the colon epithelium specific CDX2P regulatory elements with the loxP-CreER^T2^ system has enabled Apc inactivation in the adult mouse colon epithelium ^9,10^. In previous work, we showed that loss of intestinal keratin 8 (K8) makes the colon more susceptible to tumor development upon a second carcinogenic hit with AOM ^11^. In the present study, we aimed to avoid AOM toxicity and develop a genetic mouse model for colon tumorigenesis by inactivating K8 and Apc specifically in colon epithelial cells.

Keratin 8 (K8) is an integral cytoskeletal component of intestinal epithelial cells. Healthy intestinal epithelium primarily expresses type I (K18–K20) keratins paired with type II (K7–K8) keratins. However, K7 is only expressed in the mouse intestine and is undetectable in the healthy human intestine ^12,13^. Loss of the major type II keratin K8 in mice dramatically reduces the expression of its type I partners, K18 and K19, and to a lesser extent, type II K7 in the colon. This leads to a colon phenotype characterized by diarrhea, hyperproliferation, epithelial damage, modest inflammation, imbalanced differentiation, and experimentally induced tumorigenesis ^11,14,15^. One of the earliest pathological events following K8 loss is colonocyte hyperproliferation ^16^. After K8 loss, proliferative signaling activity is increased in the colonic epithelium ^11,14^.

Given the involvement of K8 in multiple protective roles in the colon, we investigated whether combining the loss of K8 and Apc in colon epithelial cells could enhance and accelerate colon tumorigenesis in mice. To drive the loss of K8 and Apc in colon epithelial cells, we used the CDX2P sequence-dependent loxP-CreER^T2^ system. Our new K8^flox/flox^; CDX2P-CreER^T2^ mouse model displays a patchy, localized loss of K8, accompanied by epithelial changes in the K8-negative areas of the colon. When combined with monoallelic Apc inactivation, these mice exhibit significantly enhanced and accelerated colon tumorigenesis compared to mice with monoallelic Apc inactivation and intact K8. Additionally, human colon adenocarcinoma patients show decreased transcriptional expression of K8 in multiple comparisons. The model we generated is rapid, genetic, and highly relevant to human CRC, as it involves Apc inactivation, reduced K8 expression, and tumor development specifically in the colon. These findings highlight the essential role of colonocyte K8 in maintaining epithelial integrity and protecting against tumorigenesis.

## Materials and Methods

### Experimental mouse models

To generate a colon specific K8 knockdown mouse model, K8^flox/flox^ mice (C57BL/6) from Prof. Karen M. Ridge (Northwestern University, Chicago Illinois, USA) ^11^ were bred with CDX2P-CreER^T2^; Apc^flox/+^ mice (C57BL/6) provided by Prof. Yatrik M. Shah (University of Michigan, Ann Arbor, USA) ^9^. The offsprings K8^flox/+^; CDX2P-CreER^T2^; Apc^flox/+^ were crossed with K8^flox/flox^. The intercrosses of K8^flox/flox^; CDX2P-CreER^T2^; Apc^flox/+^ and K8^flox/flox^; Apc^flox/+^ generated both K8^flox/flox^; CDX2P-CreER^T2^ and K8^flox/flox^; CDX2P-CreER^T2^; Apc^flox/+^. Mice were housed at the Central Animal Laboratory of University of Turku and treated according to animal licenses (ESAVI/16359/2019 and ESAVI/4498/2023) approved by the State Provincial Office of South Finland.

Mice were genotyped with PuReTaq Ready-To-Go PCR Beads (Cytvia, Buckinghamshire, UK) using primers for K8 flox: (5′-GCGTGGCTTTGGGATTTAGATTAG-3′ and 5′-CCTCCAGCCATGTTTCTTTATCTC-3′), for Cre: (5’-AGTGCGTTCGAACGCTA-GAGCCTGT-3’ and 5’-GAACCTGATGGACATGTTCAGG-3’) and for Apc flox: (Apc-P3: 5’-GTTCTGTATCATGGAAAGATAGGTGGTC-3’), (Apc-P4: 5’-CACTCAAAAC-GCTTTTGAGGGTTGATTC-3’) and (Apc-P5: 5’-GAGTACGGGGTCTCTGTCTCAG-TGAA-3’).

### Tamoxifen administration

Tamoxifen (TAM) solution was prepared by dissolving tamoxifen (Sigma-Aldrich, MO, USA) in vehicle (Sigma-Aldrich, MO, USA) and 100 mg/kg TAM was used. The vehicle solution contained only corn oil. Adult 2-3 months old K8^flox/flox^; CDX2P-CreER^T2^ and K8^flox/flox^; CDX2P-CreER^T2^; Apc^flox/+^ mice received a daily tamoxifen (+TAM) or vehicle (–TAM) dose intraperitoneally for 5 consecutive days. Mice were monitored for their body weight and stool consistency. Stool consistency was scored as normal=1, formed but soft=2, slightly loose=3 and liquid=4, as previously described ^17^. K8^flox/flox^; CDX2P-CreER^T2^ and K8^flox/flox^; CDX2P-CreER^T2^; Apc^flox/+^ mice were sacrificed 28 days and 78 days after 1^st^ TAM administration, respectively. CDX2P-CreER^T2^; Apc^flox/+^ mice were used as an additional control for K8^flox/flox^; CDX2P-CreER^T2^; Apc^flox/+^ mice. All experiments had an equal representation of males and females.

### Tissue collection and processing

Mice were euthanized by CO_2_ inhalation and whole colon was removed and washed in ice cold phosphate-buffered saline (PBS) (Medicago, Uppsala, Sweden) pH 7.4 and colon length was measured. Proximal colon (PC) and distal colon (DC) tissues were collected for histology, immunoblotting and immunohistochemistry analysis. Isolated tissues were either snap frozen and stored in liquid N_2_, fixed immediately in 4 % paraformaldehyde (PFA) (ThermoFischer Scientific, Kandel, Germany) pH 7.4, followed by paraffin embedding, or embedded in optimal cutting temperature compound (OCT) (Sakura Finetek, CA, USA) and kept at − 80 °C. Similarly, ileum tissue (approx. 2 cm distal from the caecum) was also collected for protein analysis.

### Histological examination

Paraffin embedding, processing (4 µm thick sections) and hematoxylin and eosin (H&E) staining of 4 % PFA fixed tissue samples was performed by Histocore facility at Institute of Biomedicine, University of Turku. H&E stained samples were scanned using Pannoramic 1000 slide scanner (3DHISTECH, Budapest, Hungary). Colon crypt lengths were measured and % crypt loss was analyzed using CaseViewer 2.4 (3DHISTECH, Budapest, Hungary). H&E stained individual crypts per mouse were quantified to determine mean crypt length. For each mouse, the colon mucosal regions along mucosae muscularis with prominent erosion and lack of crypts were quantitated as % crypt loss of the entire perimeter of colon ^11^.

### SDS-PAGE and Immunoblotting

Isolated total tissue samples were homogenized in ice cold homogenization buffer (0.187 M Tris-HCl pH 6.8, 3 % SDS, 5 mM EDTA) supplemented with 1 x cOmplete protease inhibitor cocktail (Roche, Mannheim, Germany) and 1 mM phenylmethylsulfonyl fluoride (PMSF) (Sigma-Aldrich, MO, USA) on ice. Afterwards, protein concentrations of each sample were quantified with Pierce bicinchoninic acid (BCA) protein assay kit (Thermo Fisher Scientific, MA, USA). Samples were prepared as 5 µg protein/10 µl using 3x Laemmli sample buffer (30 % glycerol, 3 % SDS, 0.1875 M Tris-HCl pH 6.8, 0.015 % bromophenol blue and 3 % β-mercaptoethanol). The samples were loaded on 10 % sodium dodecyl sulphate-polyacrylamide gels along with precision plus protein dual color standards (BIO-RAD, CA, USA) to determine the molecular weight of the separated proteins. Proteins were then transferred to polyvinylidene fluoride (PVDF) (Cytvia, Buckinghamshire, UK) membranes and immunoblotted for proteins of interest. The primary antibodies used were: rabbit anti-K7 (181598) from Abcam (Cambridge, UK), rat anti-K8 (Troma I) from Developmental Studies Hybridoma Bank (IA, USA), mouse anti-K18 (61028) from Progen (Heidelberg, Germany), rat anti-K19 (MABT913) from EMD Millipore Corporation (CA, USA), rabbit anti-K20 (97511) from Abcam (Cambridge, UK), sheep anti-IL22BP (AF2376) from R&D Systems (MN, USA), rabbit anti-β-actin (4967) from Cell Signaling Technology (MA, USA), mouse anti-β-tubulin (T8328) from Sigma-Aldrich (MO, USA) and rat anti-Hsc70 (SPA-815) from Enzo Life Science (NY, USA). The secondary antibodies used were: anti-rabbit Alexa-Fluor 800 (A32735) from Invitrogen (CA, USA), anti-rat Alexa-Fluor 680 (A21096) from Invitrogen (CA, USA), anti-mouse Alexa-Fluor 800 (A32730) from Invitrogen (CA, USA), anti-sheep IgG-HRP (12-342) from Sigma-Aldrich (MO, USA). The HRP signals were detected using western lightning plus-ECL (Perkin Elmer, MA, USA) and visualized with iBright CL1000 imaging system (Invitrogen, CA, USA). The fluorescent signals were visualized with iBright FL1000 imaging system (Invitrogen, CA, USA). The protein bands were quantified using ImageJ software (National Institutes of Health, MD, USA) as previously described ^18^ and normalized to their respective loading controls.

### Immunohistochemistry

Frozen OCT embedded samples were processed (6 μm thick sections) using a Leica CM 1950 Research Cryostat (Leica Microsystems, Wetzlar, Germany) and fixed with 1 % PFA in PBS (pH 7.4), for 15 minutes and afterwards immunostained as previously described ^19^. The primary antibodies used were: rabbit anti-K7 (181598) from Abcam (Cambridge, UK), rat anti-K8 (Troma I) from Developmental Studies Hybridoma Bank (IA, USA), rabbit anti-K18 (SAB4501665) from (Sigma-Aldrich, MO, USA), rat anti-K19 (MABT913) from EMD Millipore Corporation (CA, USA), rabbit anti-CDX2 (EPR2764Y) from Cell Marque (CA, USA), rabbit anti-MPO (RB-373-A0) from Thermo Fisher Scientific (MA, USA), rabbit anti-Ki67 (16667) from Abcam (Cambridge, UK) and rabbit anti-Notch-1 (sc-6014) from Santa Cruz (TX, USA). The secondary antibodies used were: anti-rabbit Alexa-Fluor 488/546 (A21206/A10040) from Invitrogen (CA, USA) and anti-rat Alexa-Fluor 488/568 (A21208/A11077) from Invitrogen (CA, USA). The nuclei were stained with DAPI (Invitrogen, CA, USA) before mounting the stained samples with ProLong Gold antifade reagent (Invitrogen, CA, USA).

Paraffin embedded samples were processed and immunolabelled for K8 (Troma I) with hematoxylin counter-stain by Histocore facility at Institute of Biomedicine, University of Turku. Imaging was done using Marianas Spinning disk confocal microscope (Intelligent Imaging Innovations, Denver, CO, USA), Pannoramic 1000 and Pannoramic MIDI slide scanners (3DHISTECH, Budapest, Hungary).

### Image analysis

ImageJ (National Institutes of Health, MD, USA) and QuPath v0.2.3 ^20^ software were used for image analysis. For overall analysis between –TAM and +TAM mice, areas inside the mucosae muscularis (towards lumen) of colon were drawn. For analysis of differences in K8-positive (K8+) and K8-negative (K8–) areas in +TAM mice, K8+ or K8– crypts were selected based on K8 immunofluorescence staining. The number of MPO+ cells were counted using the counting tool in QuPath and presented as number of MPO+ cells per mm^2^ area. Ki67 cell positivity and Notch-1 signal intensity were quantified in ImageJ as previously described ^21,22^. For Ki67 quantification, the Otsu threshold method was used and watershed separation was applied for overlapping nuclei. Total number of nuclei (based on DAPI immunofluorescence) and Ki67+ nuclei were determined using the Analyze particles tool with the following parameters (size=10-infinity, circularity=0-1) and presented as percentage of Ki67+ cells per total number of cells. For quantifying the fluorescence intensity of Notch-1, mean gray value, area and integrated intensity and their background signals were measured and shown as fold change of mean fluorescence intensity corrected against the background signal. K8 positivity (based on DAB staining) in dysplastic areas and colon adenocarcinoma was determined using QuPath positive cell detection tool as previously described ^23^.

### TCGA analysis

Transcriptional expression of K8 was determined in multiple comparisons on the COAD-TCGA dataset using UALCAN ^24,25^. K8 expression analysis was conducted between sample types (normal and primary tumor), histological subtypes (normal, adenocarcinoma and mucinous adenocarcinoma), individual cancer stages (normal, stage1, stage2, stage3 and stage4), nodal metastasis status (normal, N0, N1 and N2), patient’s age in years (normal, 21-40, 41-60, 61-80 and 81-100) and gender (normal, male and female).

### Statistical analysis and data preparation

Results were analyzed and graphs were generated using Prism 9 (GraphPad Software, San Diego, CA, USA) and figures were prepared in Adobe Illustrator 2024 (Adobe, Inc., San Jose, CA, USA). The statistical significance between two groups was determined after unpaired t test. In a comparison between two groups analyzed at multiple time points, two-way ANOVA Bonferroni’s post hoc test was utilized. The significance is indicated as * P < 0.05; ** P < 0.01; *** P < 0.001; **** P < 0.0001.

## Results

### Colon specific K8 downregulation in tamoxifen-treated K8^flox/flox^; CDX2P-CreER^T2^ mice leads to mild diarrhea and reduced expression of major colonic keratins

The colon epithelium specific targeting of K8 was achieved by crossing mice with floxed Krt8 gene ^11^ to mice expressing a tamoxifen inducible CDX2P-CreER^T2^ transgene ^9^, and thereby generating the colon specific K8^flox/flox^; CDX2P-CreER^T2^ mouse model. Experimental mice received tamoxifen (TAM) to disrupt K8 expression in colon epithelial cells and control mice received the vehicle (corn oil). Mice were monitored for their body weight and stool consistency and samples were collected on day 28 **(Fig. 1A)**. There was no difference in body weight of mice between groups until day 28 **(Fig. 1B)**. However, TAM-treated mice had on average looser stool than the vehicle-treated mice **(Fig. 1C)**. Mean colon length did not differ between TAM-treated mice and vehicle-treated mice (8.3 ± 0.5 cm versus 7.3 ± 0.2 cm, respectively p = 0.055).

**Figure 1:**
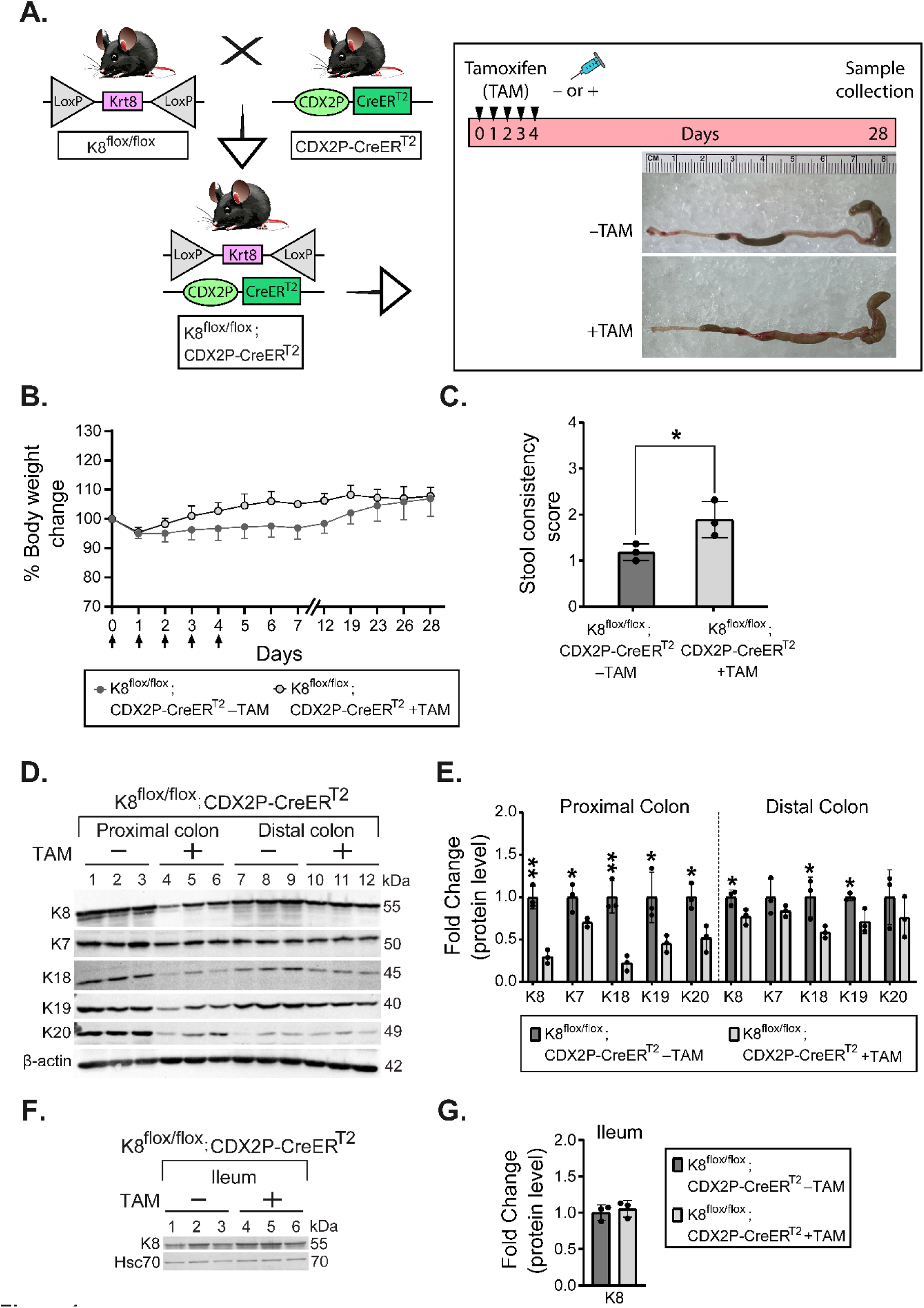
Colon epithelial cell K8 downregulation in TAM-treated K8^flox/flox^; CDX2P-CreER^T2^ mice leads to mild diarrhea and reduced protein levels of major keratins in the colon. A) Schematic representation of K8^flox/flox^; CDX2P-CreER^T2^ mouse model generation and experimental study timeline, black arrowheads indicate vehicle (–TAM) and tamoxifen (+TAM) administrations. Representative colon images on day 28. B) Percentage body weight changes of K8^flox/flox^; CDX2P-CreER^T2^ (– TAM/+TAM) mice was determined and presented as mean (n=3 mice per group) ± SD at different time points during the experimental study, black arrows indicate – TAM/+TAM administrations. C) Stool consistency of K8^flox/flox^; CDX2P-CreER^T2^ (– TAM/+TAM) mice was scored and presented as mean (n=3 mice per group, each data point represents an individual mouse) ± SD. D) Total proximal colon lysates from K8^flox/flox^; CDX2P-CreER^T2^ –TAM (Lanes 1–3), K8^flox/flox^; CDX2P-CreER^T2^ +TAM (Lanes 4–6) and total distal colon lysates from K8^flox/flox^; CDX2P-CreER^T2^ –TAM (Lanes 7–9), K8^flox/flox^; CDX2P-CreER^T2^ +TAM (Lanes 10–12) were immunoblotted for K8, K7, K18, K19 and K20. β-actin was used as the loading control. E) Western blots from D were quantified and normalized to β-actin. The results are presented as mean (n=3 mice per group, each data point represents an individual mouse) protein fold changes ± SD. F) Total ileum lysates from K8^flox/flox^; CDX2P-CreER^T2^ –TAM (Lanes 1– 3) and +TAM (Lanes 4–6) were immunoblotted for K8 and Hsc70 was used as the loading control. G) Western blot from F was quantified and normalized to Hsc70 and presented as mean (n=3 mice per group, each data point represents an individual mouse) protein fold changes ± SD. The statistical significance was determined after performing two-way ANOVA Bonferroni’s post hoc test for B and unpaired student’s t test for C, E and G, shown as * P < 0.05 and ** P < 0.01.

Proximal and distal colon tissue were analyzed for protein levels of K8, the other type II (K7) and type I keratins (K18, K19 and K20). K8 protein levels were downregulated in both proximal and distal colon, followed by reduced K18 and K19 protein expression levels. However, K7 and K20 protein levels were significantly reduced only in the proximal colon **(Fig. 1D–E)**. In the ileum, K8 protein levels remained unchanged when compared between vehicle-treated and TAM-treated mice as expected **(Fig. 1F–G)**. These findings indicate that TAM-treated K8^flox/flox^; CDX2P-CreER^T2^ mice show a significant downregulation of K8 in the colon rather than a complete K8 inactivation 28 days after 1^st^ TAM administration, with unaltered K8 protein levels in the ileum.

### CDX2P-CreER^T2^-mediated recombination in tamoxifen-treated K8^flox/flox^; CDX2P-CreER^T2^ mice results in mosaic K8 disruption with crypt loss in K8-negative areas

To investigate the observed downregulation and incomplete deletion of K8 in the colon, we analyzed K8 expression patterns in the colonic epithelium using immunofluorescence staining. K8 inactivation was partial, resulting in a patchy knockout pattern characterized by regions lacking K8 expression (K8-negative) interspersed with regions retaining K8 expression (K8-positive). Notably, the proximal colon exhibited a higher prevalence of K8-negative areas compared to the distal colon **(Fig. 2A, Supplemental Fig. 1A)**. Similarly, K7, K18, and K19 demonstrated patchy expression patterns that mirrored K8 staining in both the proximal and distal colon **(Supplemental Fig. 1B)**.

**Figure 2:**
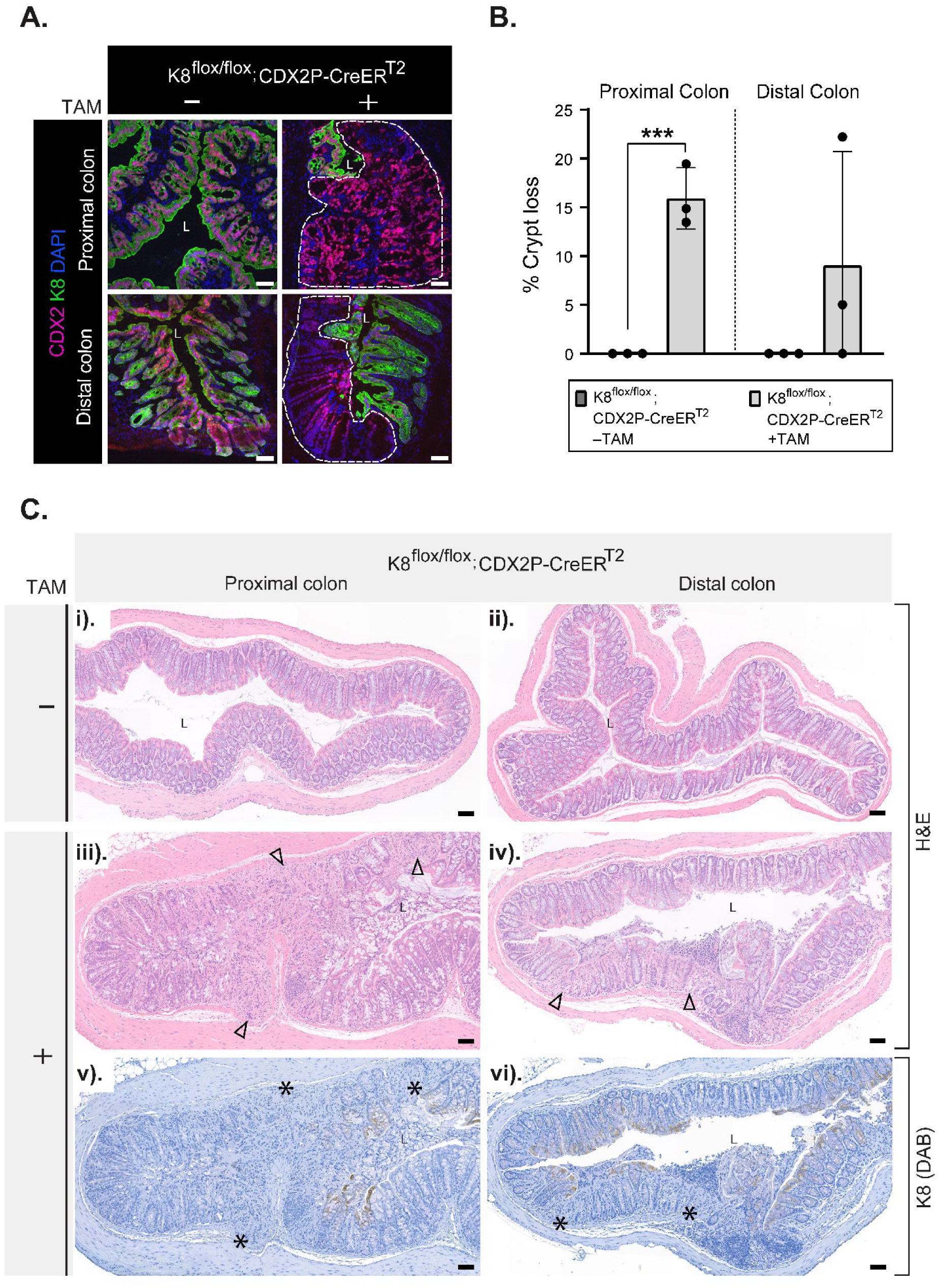
K8 downregulation in K8^flox/flox^; CDX2P-CreER^T2^ mice shows patchy knockout pattern in the colon and crypt damage occurs in K8-negative areas. **A)** Immunofluorescence staining of K8 (green), CDX2 (magenta) and nuclei, DAPI (blue) in proximal and distal colon sections of K8^flox/flox^; CDX2P-CreER^T2^ (–TAM/+TAM) mice (n=3 mice per group) is shown. Areas within the white dashed lines represent K8-negative colon crypts, L=Lumen, Scale bar=50μm. **B)** Percentage of crypt losswas determined from hematoxylin and eosin (H&E) stained proximal and distal colon sections of K8^flox/flox^; CDX2P-CreER^T2^ (–TAM/+TAM) and presented as mean (n=3 mice per group, 3 sections per proximal/distal colon) ± SD, each data point represents an individual mouse. **C)** i-iv) H&E sections from K8^flox/flox^; CDX2P-CreER^T2^ (– TAM/+TAM) mice (n=3 mice per group) with black arrowheads indicate crypt loss in the proximal and distal colon, L=Lumen, Scale bar = 100μm. v-vi) K8 (DAB) immunolabeling in the same proximal and distal colon sections of K8^flox/flox^; CDX2P-CreER^T2^ +TAM mice (n=3 mice) with black asterisks indicate K8-negative areas, L=Lumen, Scale bar = 100μm. All the images are representative of n=3 mice per group. The statistical significance was determined after performing unpaired student’s t test for B, shown as * P < 0.05, ** P < 0.01 and *** P < 0.001.

Next the histological phenotype in the colon of TAM-treated K8^flox/flox^; CDX2P-CreER^T2^ mice was assessed, focusing on crypt damage **(Fig. 2C)**. Crypt loss was quantified separately in the proximal and distal colon. Mice with K8 downregulation exhibited significantly higher crypt loss in the proximal colon compared to the distal colon, where substantial inter-mouse variability was observed **(Fig. 2B)**. Importantly, crypt loss was confined to K8-negative regions of the colon **(Fig. 2C iii–vi)**. Collectively, these findings indicate that K8 downregulation is more pronounced in the proximal colon, which correlates with increased crypt loss, whereas the distal colon exhibits fewer K8-negative areas and less crypt damage.

### K8-negative areas in the proximal colon of tamoxifen-treated K8^flox/flox^; CDX2P-CreER^T2^ mice exhibit increased number of myeloperoxidase (MPO)+ cells

Since crypt loss was localized to K8-negative areas in these mice, we investigated whether colitis develops and whether K8-negative regions promote neutrophil recruitment. Myeloperoxidase (MPO)-positive cells were quantified in the lamina propria, spanning from the lumen to the muscularis mucosa, in both the proximal and distal colon. Tamoxifen-treated mice exhibited a significant increase in MPO+ cells in the proximal colon compared to vehicle-treated controls, while the distal colon showed minimal MPO+ cell infiltration, with no significant difference between treatment groups **(Fig. 3A–B)**. Within the proximal colon of TAM-treated mice, K8-negative areas exhibited significantly higher MPO+ cell infiltration compared to K8-positive areas in the same tissue **(Fig. 3C)**. In contrast, no differences in MPO+ cell numbers were observed between K8-negative and K8-positive areas in the distal colon **(Fig. 3C)**. These findings indicate that neutrophil recruitment is selectively increased in K8-negative regions of the proximal colon, suggesting that K8 deficiency drives localized colonic inflammation in this model.

**Figure 3:**
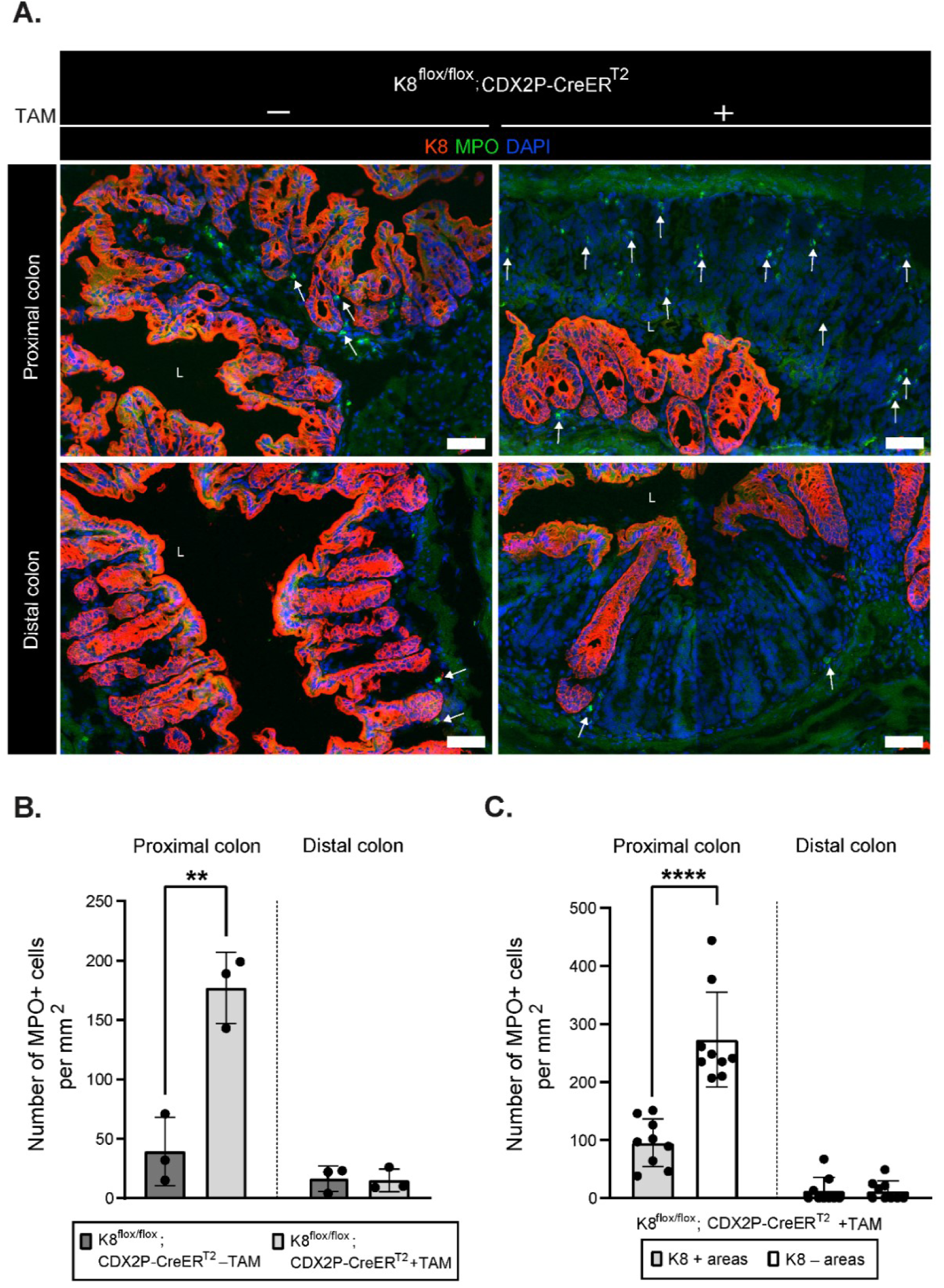
TAM-treated K8^flox/flox^; CDX2P-CreER^T2^ mice have more MPO+ cells in K8-negative areas of the proximal colon. **A)** Immunofluorescence staining of K8 (red), MPO (green), nuclei, DAPI (blue) in proximal and distal colon section of K8^flox/flox^; CDX2P-CreER^T2^ (–TAM/+TAM) mice (n=3 mice per group) is shown, white arrows indicate MPO+ cells, L=Lumen, Scale bar=100μm. Images are representative of n=3 mice per group. **B)** MPO+ cells within proximal and distal colon mucosal area were quantified from K8^flox/flox^; CDX2P-CreER^T2^ (–TAM/+TAM) mice and presented as mean (n=3 mice per group, 1 section per proximal/distal colon) ± SD, each data point represents an individual mouse. **C)** MPO+ cells within K8+ and K8– areas of proximal and distal colon were counted in K8^flox/flox^; CDX2P-CreER^T2^ +TAM mice and results are presented as mean (n=3 mice, 3 K8+ and 3 K8– areas per proximal/distal colon) ± SD, each data point represents an individual area. The statistical significance was determined after performing unpaired student’s t test for B and C, shown as * P < 0.05, ** P < 0.01, *** P < 0.001 and **** P < 0.001.

### K8-negative areas in the colon of tamoxifen-treated K8^flox/flox^; CDX2P-CreER^T2^ mice have increased number of dividing Ki67+ cells

The differences in crypt length between experimental groups, as well as between crypts with and without K8 expression in TAM-treated K8^flox/flox^; CDX2P-CreER^T2^ mice, were analyzed. In TAM-treated mice, crypt lengths in the proximal colon were significantly increased compared to vehicle-treated controls. In contrast, no statistically significant differences were observed in crypt lengths in the distal colon compared to vehicle-treated mice with intact K8 **(Fig. 4A)**. Within TAM-treated mice, crypts lacking K8 in the proximal and distal colon showed no difference in length compared to crypts retaining K8 expression **(Fig. 4B)**. To assess the proliferative activity of colonocytes in these mice, the percentage of Ki67+ cells was quantified. Overall, both the proximal and distal colons of TAM-treated mice exhibited a higher percentage of Ki67+ cells compared to vehicle-treated controls **(Fig. 4C–D)**. Notably, K8-negative crypts contained a significantly higher percentage of Ki67+ cells in both the proximal and distal colon compared to K8-positive crypts in TAM-treated K8^flox/flox^; CDX2P-CreER^T2^ mice **(Fig. 4E)**. Interestingly, K8-positive crypts in the proximal colon of TAM-treated mice were significantly taller and contained more Ki67+ cells compared to crypts from vehicle-treated mice with intact K8 (**Supplemental Fig. 1C–D**). These findings indicate that the loss of K8 is associated with increased colonocyte proliferation in crypts, correlating with the observed increase in crypt length in the proximal colon. The data also suggest that crypt proliferation and length regulation may be influenced not only by the local crypt microenvironment but also by interactions with neighboring crypts.

**Figure 4:**
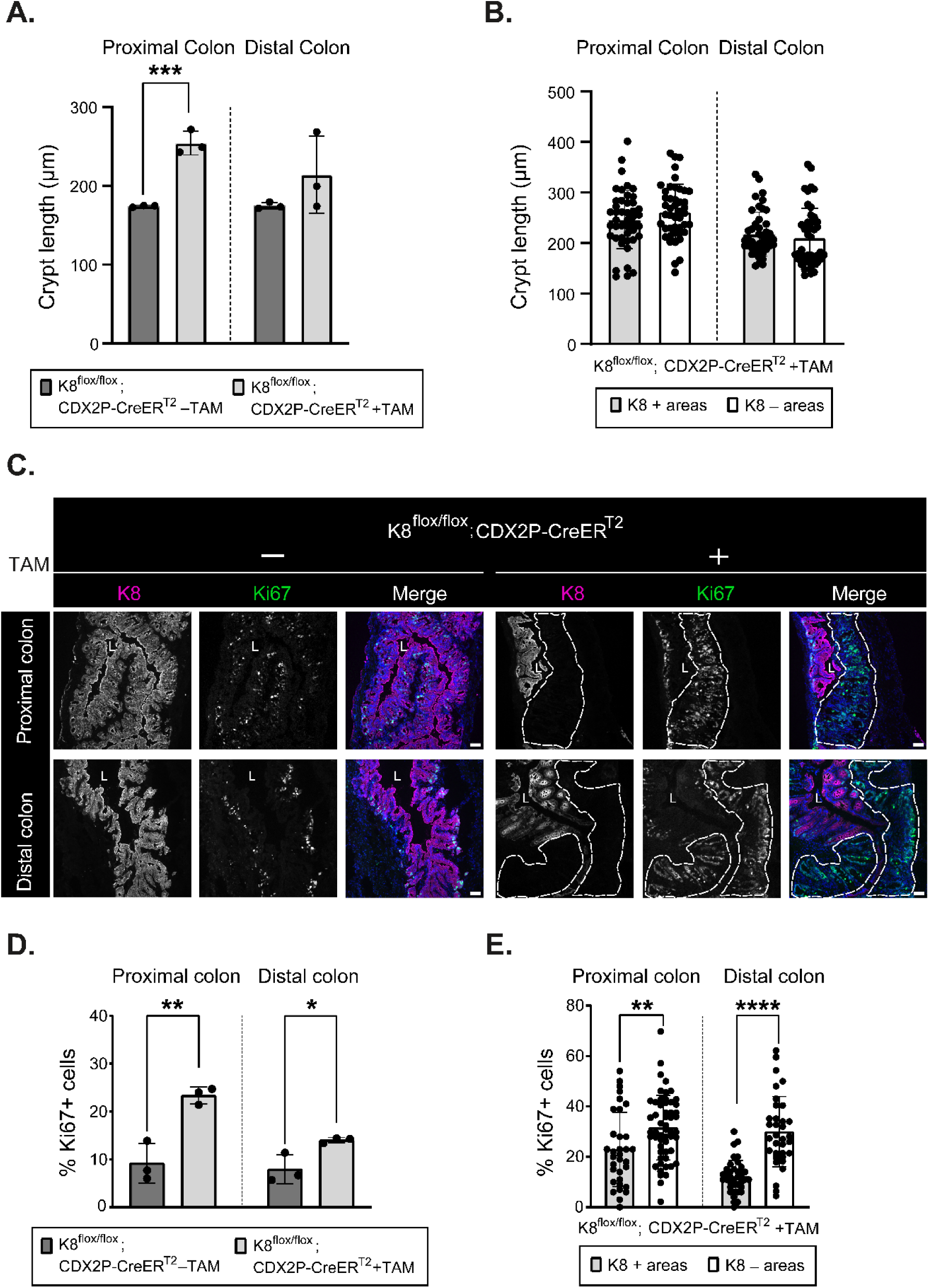
Colonic K8 downregulation based K8^flox/flox^; CDX2P-CreER^T2^ mice have lengthened crypts and more Ki67+ cells in the colon. **A)** Crypt lengths from proximal and distal colon of K8^flox/flox^; CDX2P-CreER^T2^ (–TAM/+TAM) mice were measured and presented as mean (n=3 mice per group, 30 crypts per proximal/distal colon) ± SD, each data point represents an individual mouse. **B)** Lengths of K8+ and K8– crypts from proximal and distal colon of K8^flox/flox^; CDX2P-CreER^T2^ +TAM mice were quantified and presented as mean (n=3 mice, 15 K8+ and 15 K8– crypts per proximal/distal colon) ± SD, each data point represents an individual crypt. **C)** Immunofluorescence staining of K8 (magenta), Ki67 (green), nuclei, DAPI (blue) in proximal and distal colon sections of K8^flox/flox^; CDX2P-CreER^T2^ (–TAM/+TAM) mice (n=3 mice per group) is shown. Areas within the white dashed lines represent K8-negative colon crypts, L=Lumen, Scale bar=50μm. Images are representative of n=3 mice per group. **D)** Percentage of Ki67+ cells were quantified in proximal and distal colon of K8^flox/flox^; CDX2P-CreER^T2^ (–TAM/+TAM) mice and presented as mean (n=3 mice per group, 3 images per proximal/distal colon) ± SD, each data point represents an individual mouse. **E)** Percentage of Ki67+ cells in K8+ and K8– crypts of proximal and distal colon was calculated in K8^flox/flox^; CDX2P-CreER^T2^ +TAM mice and presented as mean (n=3 mice, 6–15 K8+ and 8–15 K8– crypts per proximal/distal colon) ± SD, each data point represents an individual crypt. The statistical significance was determined after performing unpaired student’s t test for A–B and D–E, shown as * P < 0.05, ** P < 0.01, *** P < 0.001 and **** P < 0.001.

We next analyzed protein levels of interleukin-22 binding protein (IL-22BP) in the colons of these mice. IL-22BP expression is typically downregulated during intestinal damage, which increases IL-22 bioavailability to support epithelial proliferation and tissue repair ^26^. Following colon-specific K8 downregulation in TAM-treated K8^flox/flox^; CDX2P-CreER^T2^ mice, IL-22BP protein levels were significantly decreased in the proximal and distal colon in comparison to vehicle-treated mice with K8 **(Supplemental Fig. 2A–B)**. These findings suggest that K8 is physiologically essential for maintaining the balance of normal colonocyte proliferation.

Finally, we investigated whether Notch-1 expression differed between K8-negative and K8-positive crypts, as Notch-1 is a key regulator of colonic epithelial cell differentiation and has been previously shown to interact with K8, influencing its role in epithelial differentiation ^15^. On average, Notch-1 expression did not change significantly between TAM-treated K8^flox/flox^; CDX2P-CreER^T2^ and vehicle-treated mice **(Supplemental Fig. 2C–D).** However, within TAM-treated mice, K8-negative crypts in both the proximal and distal colon exhibited reduced Notch-1 expression compared to K8-positive crypts **(Supplemental Fig. 2E)**.

### Colonic K8 downregulation combined with monoallelic Apc inactivation renders K8^flox/flox^; CDX2P-CreER^T2^; Apc^flox/+^ mice more susceptible to colon tumorigenesis

To determine whether colonocyte K8 downregulation combined with Apc inactivation promotes colon tumor development, CDX2P-CreER^T2^; Apc^flox/+^ and K8^flox/flox^; CDX2P-CreER^T2^; Apc^flox/+^ mice were treated with tamoxifen intraperitoneally for 5 consecutive days and followed for 78 days **(Fig. 5A)**. The percentage of body weight change between TAM-treated CDX2P-CreER^T2^; Apc^flox/+^ mice and K8^flox/flox^; CDX2P-CreER^T2^; Apc^flox/+^ mice showed no difference over the course of the experiment **(Fig. 5B)**. Out of 3 TAM-treated CDX2P-CreER^T2^; Apc^flox/+^ mice, one developed a single tumor in the distal colon. In contrast, all TAM-treated K8^flox/flox^; CDX2P-CreER^T2^; Apc^flox/+^ mice had multiple tumors in the distal colon **(Fig. 5C)**. To evaluate whether these tumors originated from K8-negative areas, all tumors from both groups were analyzed for K8 expression. The single tumor in TAM-treated CDX2P-CreER^T2^; Apc^flox/+^ mice showed decreased K8 expression compared to adjacent normal colon epithelium. Conversely, all dysplastic lesions and adenocarcinomas in TAM-treated K8^flox/flox^; CDX2P-CreER^T2^; Apc^flox/+^ mice were K8-negative with patchy K8 distribution in the adjacent colonic epithelium **(Fig. 5D, Supplemental Fig. 3A–B)**.

**Figure 5:**
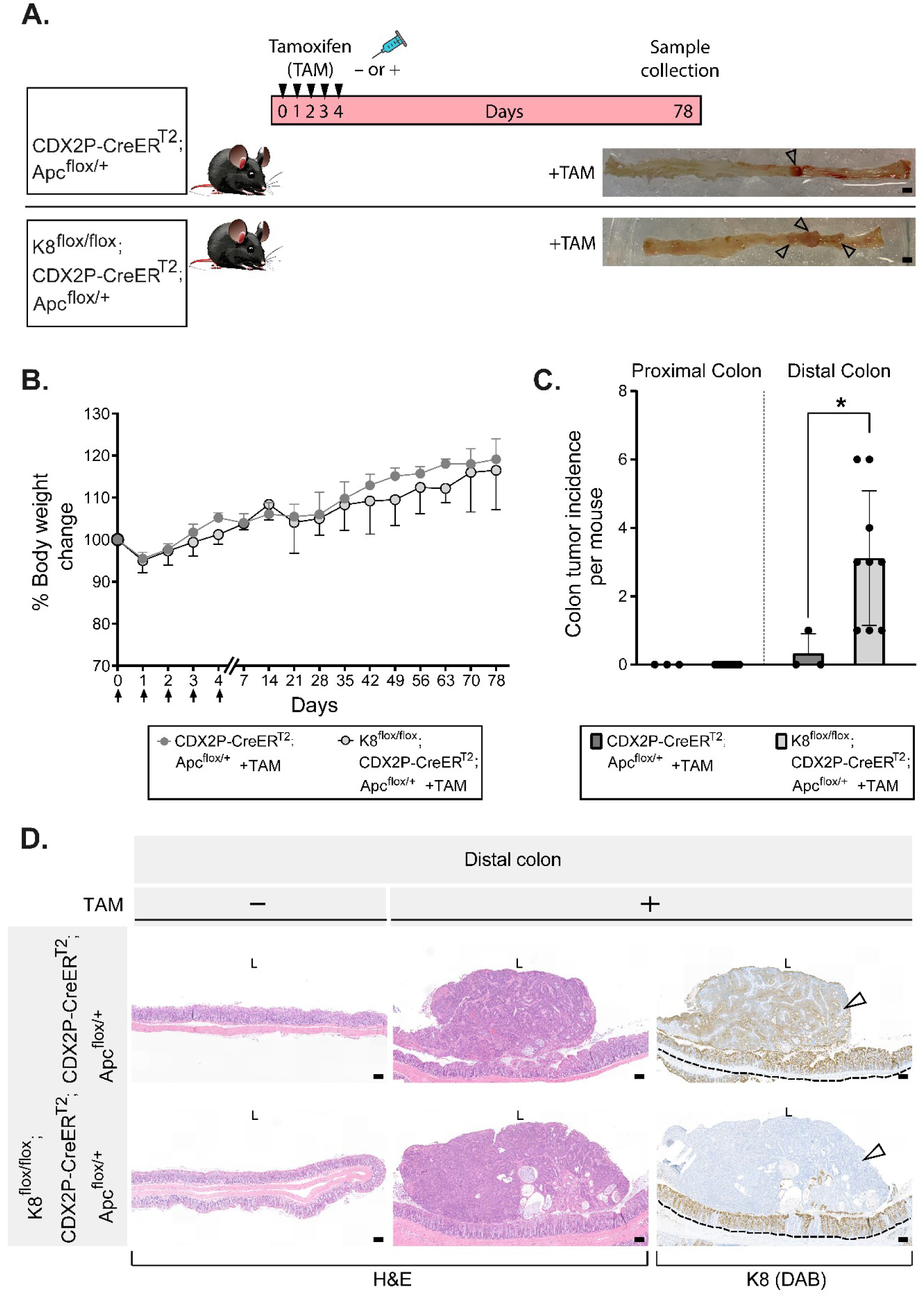
K8 downregulation in TAM-induced K8^flox/flox^; CDX2P-CreER^T2^; Apc^flox/+^ mice enhances tumor susceptibility in the distal colon. **A)** Experimental timeline for CDX2P-CreER^T2^; Apc^flox/+^ and K8^flox/flox^; CDX2P-CreER^T2^; Apc^flox/+^ mice CRC susceptibility study, black arrowheads indicate vehicle (–TAM) and tamoxifen (+TAM) administrations. Open colon images on day 78 with black arrowheads pointing toward tumors, Scale bar=5mm **B)** Percentage body weight changes of CDX2P-CreER^T2^; Apc^flox/+^ +TAM and K8^flox/flox^; CDX2P-CreER^T2^; Apc^flox/+^ +TAM mice was determined and results are shown as mean (n=3-4 mice per group) ± SD at different time points during the experimental study, black arrows indicate –TAM/+TAM administrations. **C)** Number of colonic tumors observed in CDX2P-CreER^T2^; Apc^flox/+^ +TAM (n=3) and K8^flox/flox^; CDX2P-CreER^T2^; Apc^flox/+^ +TAM (n=9), results are presented as mean ± SD, each data point represents an individual mouse. CDX2P-CreER^T2^; Apc^flox/+^ (n=3) and K8^flox/flox^; CDX2P-CreER^T2^; Apc^flox/+^ (n=9) had their own –TAM controls. **D)** H&E sections of distal colon from CDX2P-CreER^T2^; Apc^flox/+^ (–TAM/+TAM) and K8^flox/flox^; CDX2P-CreER^T2^; Apc^flox/+^ (–TAM/+TAM) and K8 (DAB) immunolabeling of colon tumors with adjacent colon. For CDX2P-CreER^T2^; Apc^flox/+^ +TAM colon tumor, black arrowhead indicates less K8 expression when compared to nearby colon tissue indicated with the black dashed line and for K8^flox/flox^; CDX2P-CreER^T2^; Apc^flox/+^ +TAM colon tumor, black arrowhead indicates negative K8 expression, with patchy K8 distribution in nearby colon tissue shown with the black dashed line, L=Lumen, Scale bar=100μm. The statistical significance was determined after performing two-way ANOVA Bonferroni’s post hoc test for B and unpaired student’s t test for C, shown as * P < 0.05.

To further characterize keratin expression, protein levels of K8 and other major colonic keratins were quantified in the colons of vehicle- and TAM-treated K8^flox/flox^; CDX2P-CreER^T2^; Apc^flox/+^ mice. K8 and other keratins were significantly downregulated in both the proximal and distal colons of TAM-treated mice compared to vehicle-treated controls **(Supplemental Fig. 4A–C)**. Interestingly, the colon tumors of TAM-treated K8^flox/flox^; CDX2P-CreER^T2^; Apc^flox/+^ mice exhibited minimal CDX2 expression compared to surrounding colonic tissue **(Supplemental Fig. 4D)**. CDX2, an intestine-specific transcription factor, controls proliferation and differentiation in the intestine epithelia ^27^. Additionally, IL-22BP protein levels were significantly reduced in both the proximal and distal colons of TAM-treated mice compared to vehicle-treated controls **(Supplemental Fig. 4E–F)**. Collectively, these findings demonstrate that K8 downregulation in the TAM-treated K8^flox/flox^; CDX2P-CreER^T2^; Apc^flox/+^ mice dramatically increases their susceptibility towards colon tumorigenesis.

### K8 expression is transcriptionally downregulated in human colon adenocarcinoma

To determine whether the correlation between K8 loss and tumorigenesis observed in mice is recapitulated in humans, K8 transcriptional expression levels were analyzed in colon adenocarcinoma and adjacent normal tissue using The Cancer Genome Atlas Colon Adenocarcinoma (TCGA-COAD) dataset mined in UALCAN. A significant reduction in K8 expression was observed in primary tumors from COAD patients, including adenocarcinoma and mucinous adenocarcinoma, compared to adjacent normal colon tissue **(Fig. 6A–B)**. Interestingly, decreased K8 expression was evident at early stages of colon cancer and was independent of metastasis. K8 expression remained reduced in patients with advanced-stage disease, including those with metastasis to 10 or more lymph nodes **(Fig. 6C–D)**. This downregulated K8 expression signature was consistent across both younger and older patients, regardless of gender **(Fig. 6E–F)**. Taken together, these findings suggest that K8 downregulation may represent an early event in the development of colon cancer.

**Figure 6:**
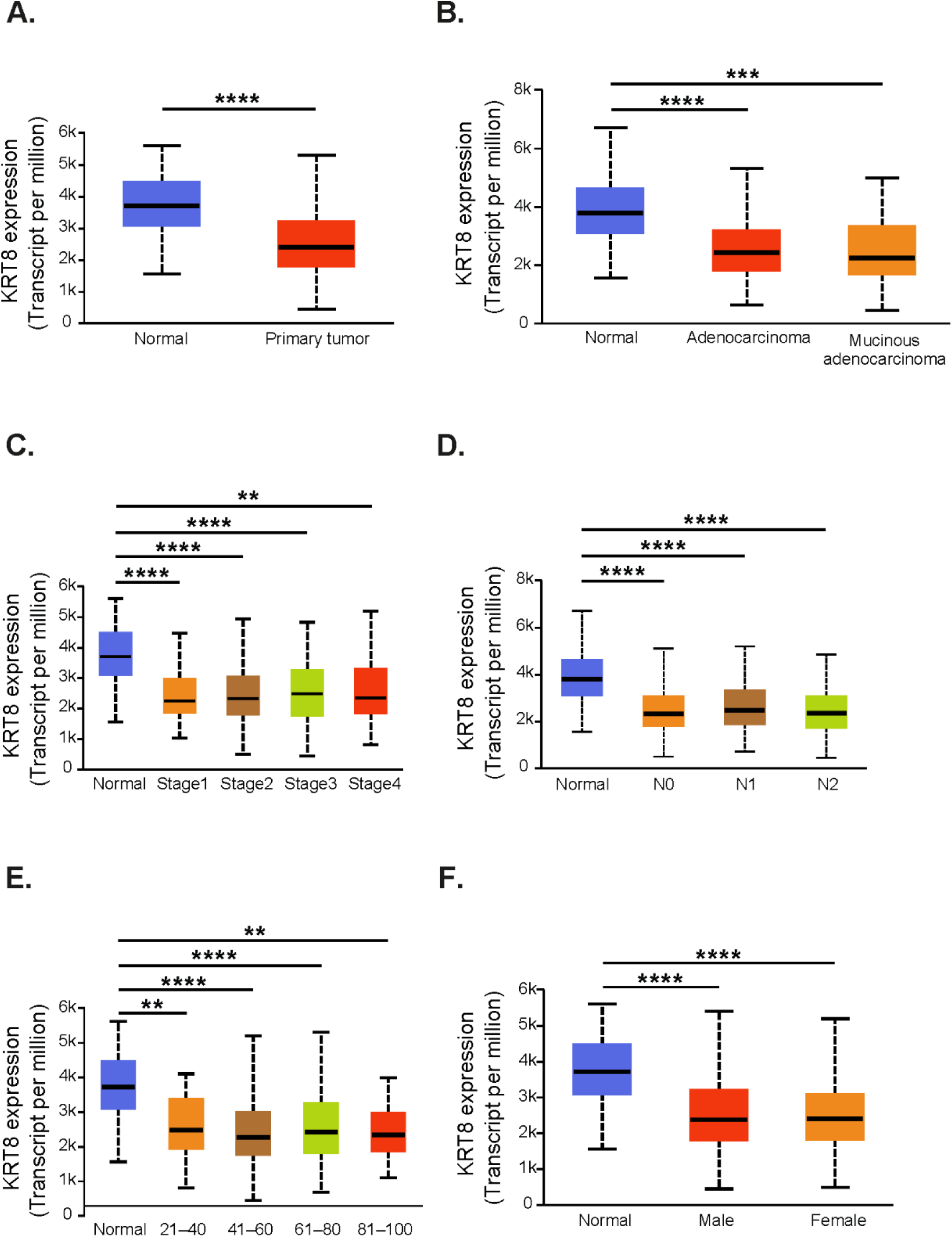
Transcriptional expression analysis using UALCAN platform reveals downregulated K8 expression in human colon adenocarcinoma in multiple comparisons. **A–F)** Comparison of K8 transcriptional expression between **A)** sample types: normal (n=41) and primary tumor (n=286) **B)** histological subtypes: normal (n=41), adenocarcinoma (n=243) and mucinous adenocarcinoma (n=37) **C)** individual cancer stages: normal (n=41), Stage1 (n=45), Stage2 (n=110), Stage3 (n=80) and Stage4 (n=39) **D)** nodal metastasis status: normal (n=41), N0 (n=166), N1 (n=70), N2 (n=47), N0: No regional lymph node metastasis, N1: Metastases in 1 to 3 axillary lymph nodes, N2: Metastases in 4 to 9 axillary lymph nodes, N3: Metastases in 10 or more axillary lymph nodes **E)** patient’s age: normal (n=41), 21–40 years (n=12), 41– 60 years (n=90), 61–80 years (n=149) and 81–100 years (n=32) **F)** patient’s gender: normal (n=41), male (n=156) and female (n=127). The statistical significance shown as * P < 0.05, ** P < 0.01, *** P < 0.001 and **** P < 0.001.

## Discussion

In this study, we report the development and phenotypic characterization of a novel mouse model for colon epithelial cell tumorigenesis driven by the combined loss of K8 and Apc (summarized in **Fig. 7**). This genetically induced model does not rely on carcinogens such as AOM, which have variable dose efficacies and long latency periods depending on the mouse strain ^28^. Mice rapidly developed colon adenocarcinoma within 78 days following the first TAM dose, exhibiting enhanced colon tumorigenesis compared to mice with Apc loss alone. The model closely mimics human CRC, which frequently features Apc inactivation and reduced K8 levels. Tumors in this model develop predominantly in the distal colon, consistent with the primary tumor sites in human CRC.

**Figure 7:**
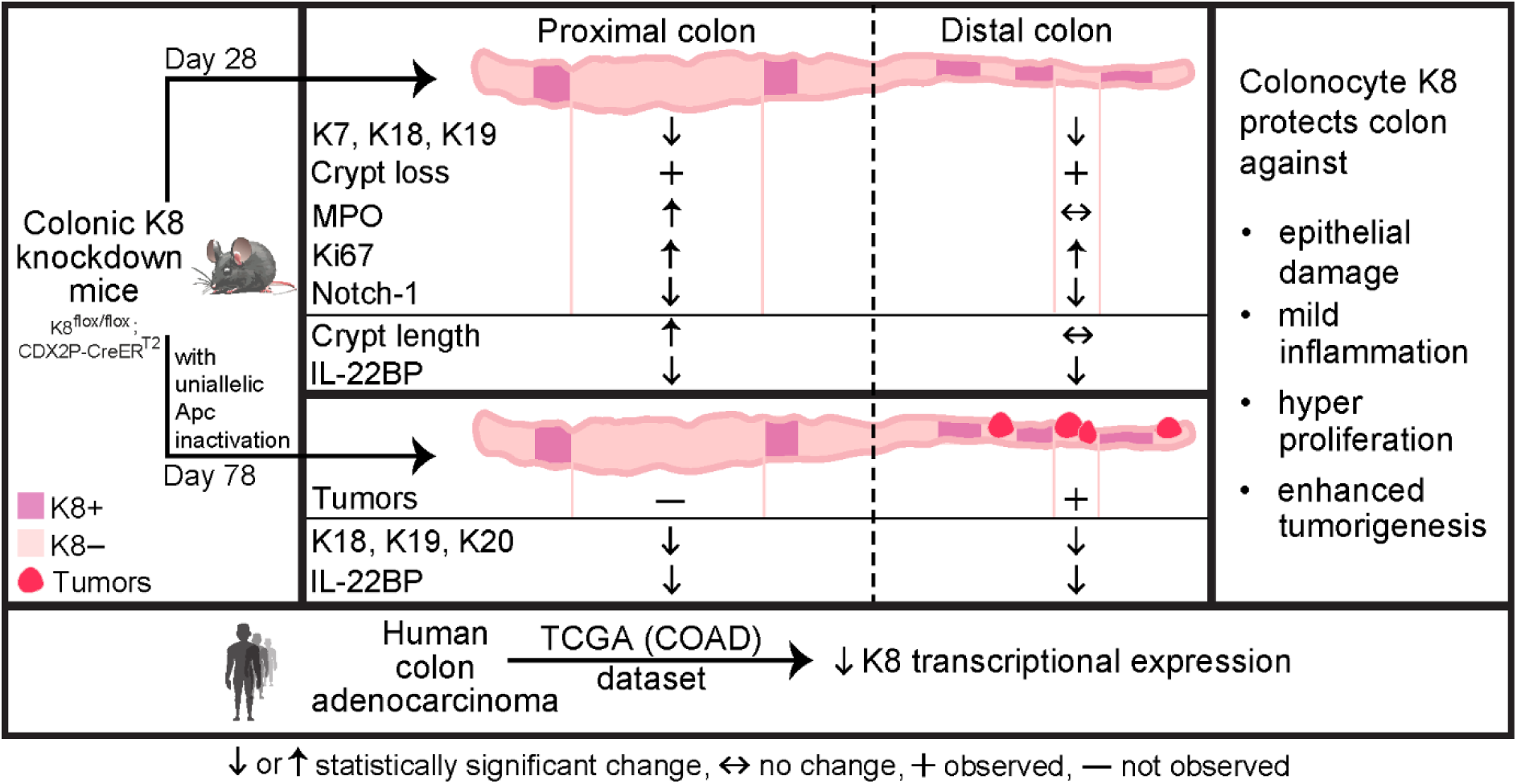
Graphical summary. Colon-specific epithelial cell tumorigenesis model. The colon-specific K8 knockdown mice display epithelial damage, crypt loss, mild inflammation (marked by increased MPO+ cells) in the proximal colon, hyperproliferation (marked by increased Ki67+ cells, overall decreased IL-22BP levels), decreased Notch-1 expression, high colorectal tumor load with monoallelic Apc inactivation, in K8-negative colonic areas. Human colon adenocarcinoma patients show decreased K8 transcriptional expression. Taken together, these local changes with colorectal tumors in the K8-negative crypts as well as recapitulating decreased K8 expression as observed in human CRC make this model resemble human CRC. It also highlights colon tissue specific role of colonocyte K8 in maintaining colon epithelial barrier integrity and protecting against colon tumorigenesis.

Existing Apc-based mouse models (Apc^1638/+^, Apc**^Δ^**^716/+^, Apc**^Δ^**^14/+^, Apc^Min-FCCC/+^) typically result in a high tumor burden in the small intestine rather than the colon ^6,29–32^. Moreover, models such as Apc^Min/+^, Apc**^Δ^**^14/+^, Apc^Min-FCCC/+^ also develop tumors in the mammary gland ^31–33^. CDX2P-NLS Cre; Apc^+/loxP^ mice have shifted tumor load towards colon, with still some tumors in the small intestine, whereas Villin-Cre; Apc^+/loxP^ mice predominantly develop tumors in the small intestine ^34^. By incorporating TAM-regulated CreER^T2^ under the CDX2P promoter, biallelic Apc inactivation was targeted to the colonic epithelium in adult CDX2P-CreER^T2^; Apc^flox/flox^ mice ^9^. Our model offers a significant advantage over these Apc mutant models, as tumors are restricted to the distal colon. Other multigene-based CRC models often involve prolonged endpoints, lethal phenotypes, and complex breeding strategies ^35^. These models do not acquire the mutations sequentially but are induced at the same time. Additionally, all the cells contain the mutation of interest, which is not the case in human CRC ^36^. Most CRC cases (75-80%) arise as a result of sequential accumulation of somatic gene alterations over time ^37^.

The complete loss of K8 throughout the intestinal epithelium of K8^flox/flox^; Villin-Cre and K8^flox/flox^; Villin-CreER^t2^ mice provided a valuable tool to study the tissue-protective roles of keratins and the sequence of pathological events following K8 deletion ^11,16^. Our new genetic model, however, not only circumvented the AOM-related challenges but importantly, K8 levels remained unchanged in the ileum, making the patchy knockout pattern specific to the colon epithelium. This patchiness mirrors the distribution of K8 expression in human CRC, where undifferentiated tumors show patchy K8 expression, while differentiated tumors exhibit stronger expression at the crypt top than at the crypt bottom ^38^.

A mild colitis-like phenotype in TAM-treated K8^flox/flox^; CDX2P-CreER^T2^ mice highlights the role of basal keratin levels in maintaining epithelial integrity. Reduced keratin levels have been associated with compromised stress-protective functions, as seen in K8^+/–^ mice, which exhibit moderate colon hyperproliferation and altered ion transport without overt inflammation ^17,39^. However, heterozygous K8 deletion increases susceptibility to dextran sulfate sodium (DSS)-induced inflammation and AOM/DSS- or Apc^Min/+^/DSS-induced CRC ^14,17,40^.

The colon epithelial damage and modest inflammation in TAM-treated K8^flox/flox^; CDX2P-CreER^T2^ mice occurred in a K8 dependent manner. However, the modest inflammation occurred only in the proximal colon of these mice. One potential cause can be different immune cell populations in proximal and distal colon ^41–43^. In TAM-treated K8^flox/flox^; CDX2P-CreER^T2^ mice, K8-negative colonic crypts exhibited increased proliferation and decreased IL-22BP protein levels, creating a pro-proliferative environment. This aligns with previous findings linking early K8 loss to the activation of proliferative pathways, including pRb, IL-22BP, and STAT3 ^11,14,16^. IL-22BP deficiency has been associated with prolonged epithelial proliferation and increased susceptibility to colon tumorigenesis in AOM/DSS and Apc^Min/+^ models ^26^. In a primary CRC patients dataset, IL-22BP was also downregulated ^44^. IL-22 treatment *in vivo* and *in vitro* increased intestinal epithelial cell proliferation by inhibiting Notch-1 and Wnt signaling ^45^. Notch-1 expression was also slightly decreased in K8-negative colonic crypts of our TAM-treated mice. These results altogether, suggest that the colon epithelial K8 is essential for normal proliferation activity of colonocytes and protects the colonocyte against epithelial hyperproliferation and colon tumorigenesis. However, the direct causative mechanism behind it needs to be elucidated.

Interestingly, despite modest inflammation and proliferation in the proximal colon of our model, no tumors were observed in this region. This may be due to increased crypt loss and colonocyte detachment in the proximal colon. Tumor development primarily in the distal colon suggests that malignant colonocytes initially require a keratin cytoskeleton for attachment and tumor formation. This anatomical tumor distribution makes our model particularly relevant for studying left-sided CRC, which often initiates with Apc inactivation ^46–48^. In our model, the precise loss of K8 and Apc accelerates tumorigenesis in the distal colon before malignant colonocytes detach, consistent with the “just-right” hypothesis, which suggests that Apc inactivation and downstream signaling must occur within an optimal range to drive tumor initiation ^49,50^. Importantly, no metastasis was observed within 78 days, indicating that this model faithfully replicates primary colon tumorigenesis. As such, it represents a valuable tool for preclinical studies aimed at evaluating therapeutic responses in CRC.

Decreased K8 expression in human colon adenocarcinoma, as observed in the UALCAN database analysis, further supports the protective role of K8 in the colon. Consistent with this, one of the control TAM-treated CDX2P-CreER^T2^; Apc^flox/+^ mice in our study, which retained K8 expression and developed a single colon tumor, exhibited lower K8 levels in the tumor compared to the adjacent colon tissue. A reduction in K8 expression in CRC tissue relative to adjacent normal colon has also been reported ^40^. Interestingly, multiple K8 isoforms were found to be elevated in the adjacent normal colon tissue of patients with colon polyps or tumors, suggesting that K8 expression may increase in adjacent normal mucosa as adenomas progress to carcinomas ^51^. This finding aligns with our model, where the colon shows a distribution of K8-negative crypts adjacent to K8-positive crypts, potentially reflecting the diseased and normal epithelium seen in human CRC. However, a more detailed analysis of K8 expression differences between healthy colon tissue, morphologically normal adjacent colon mucosa, and CRC tissues is needed. Most studies and databases typically compare paired CRC tumors with adjacent normal tissue from the same patient, but it is critical to assess K8 expression across a broader spectrum—from healthy colon mucosa to morphologically normal adjacent tissue and CRC lesions. This is particularly important because morphologically normal tissue near a polyp or tumor may be pre-cancerous and not truly representative of healthy colon tissue. Our data also suggests a lateral signaling effect from K8-negative crypts to neighboring K8-positive crypts, as the K8-positive crypts in TAM-treated mice were significantly taller and had more Ki67+ cells compared to vehicle-treated controls.

In the UALCAN database analysis, K8 expression was already reduced at stage 1 in patients without nodal metastasis, suggesting that K8 downregulation is an early event in CRC development. The loss of K8/K20 expression has been implicated in the epithelial-to-mesenchymal transition (EMT), with low K8/K20 expression correlating with poor survival in CRC patients due to the aggressive nature of their disease ^52^. Interestingly, the colon tumors in our model were CDX2-negative, despite CDX2 expression in the adjacent colon epithelium. Loss of CDX2 is associated with more aggressive CRC and increased EMT ^53^. The K7/K20/CDX2 multi-marker is commonly used to assess metastatic CRC and is found in microsatellite stable tumors ^54^. It would be valuable to investigate K8 expression in different CRC subtypes and stages. Moreover, the relationship between K8 and CDX2 expression should be explored to assess their potential role in specific CRC subtypes.

In conclusion, we present an improved colon-specific epithelial cell tumorigenesis mouse model with the following key features: (i) a local deficiency of K8 in colon epithelial cells, resulting in a mild colitis-like phenotype with epithelial damage, increased inflammation, and a pro-proliferative crypt environment; (ii) an increased number of tumors in the colon epithelium when K8 downregulation is combined with monoallelic Apc loss, with (iii) no effect on K8 levels in other organs; (iv) a patchy, localized loss of K8 in the colon epithelium that mirrors the local changes observed in human CRC; and (v) a model that recapitulates the reduction of K8 expression seen in human colorectal tumors. These features make this model a promising candidate for preclinical CRC research, including drug testing. Importantly, this study enhances our understanding of the essential role of K8 in colonocyte protection of the epithelial barrier and its contribution to tumor suppression in the colon.

## Acknowledgements

We are grateful to Professor Karen M. Ridge (Northwestern University, USA) for the donation of the transgenic K8^flox/flox^ mouse strain. We thank the members of the Toivola Epithelial Biology Lab (Åbo Akademi University), especially Jimmy Fagersund and Theresia Jansson for their assistance in initial mice screening, and Professor Pekka Taimen (University of Turku) for advice on tumor histopathology. Imaging was performed at The Cell Imaging and Cytometry (CIC) Core, Turku Bioscience Center (University of Turku and Åbo Akademi University), Biocenter Finland, Turku, Finland. Histological methods were performed by Histology core facility of the Institute of Biomedicine, University of Turku, Finland. This project was supported by Academy of Finland project Grants 315139, 332582 including InFLAMES Flagship Programme, 337531 357911; Åbo Akademi University Center of Excellence in Cellular Mechanostasis, and Solutions for Health; Medicinska understödsföreningen Liv och Hälsa foundation (to D.M.T.); NovoNordisk Foundation (NNF23OC0087039) to (L.P.); Suomen Kulttuurirahasto, Varsinais-Suomi Regional Fund (to M.T.); K. Albin Johansson Foundation (to M.T. and C.-G.A.S.); Svenska kulturfonden (to M.T., M.M.E.M. and C.-G.A.S.); Victoriastiftelsen (to C.-G.A.S.); Makarna Olins Fund (to C.-G.A.S.); Ida Montinin säätiö (to M.T.); Åbo Akademi University Foundation (to M.M.E.M.); Doctoral Network Molecular Biosciences (to M.T. and C.-G.A.S.); and NCI R01CA148828, R01CA245546, and NIDDK R01DK095201 grants (to Y.M.S).

## Author contributions

M.T., C.-G.A.S., Y.M.S. and D.M.T. conceived and designed research; M.T., M.M.E.M., C.-G.A.S., L.P., and D.M.T. performed experiments; M.T., M.M.E.M., L.P., and D.M.T. analyzed data; M.T. prepared the figures; M.T. with contributions from L.P., and D.M.T. drafted manuscript; M.T., M.M.E.M., C.-G.A.S., L.P., Y.M.S., and D.M.T. edited and revised manuscript; M.T., M.M.E.M., C.-G.A.S., L.P., Y.M.S., and D.M.T. approved final version of manuscript.

## Supplemental figure

**Supplemental Figure 1:**
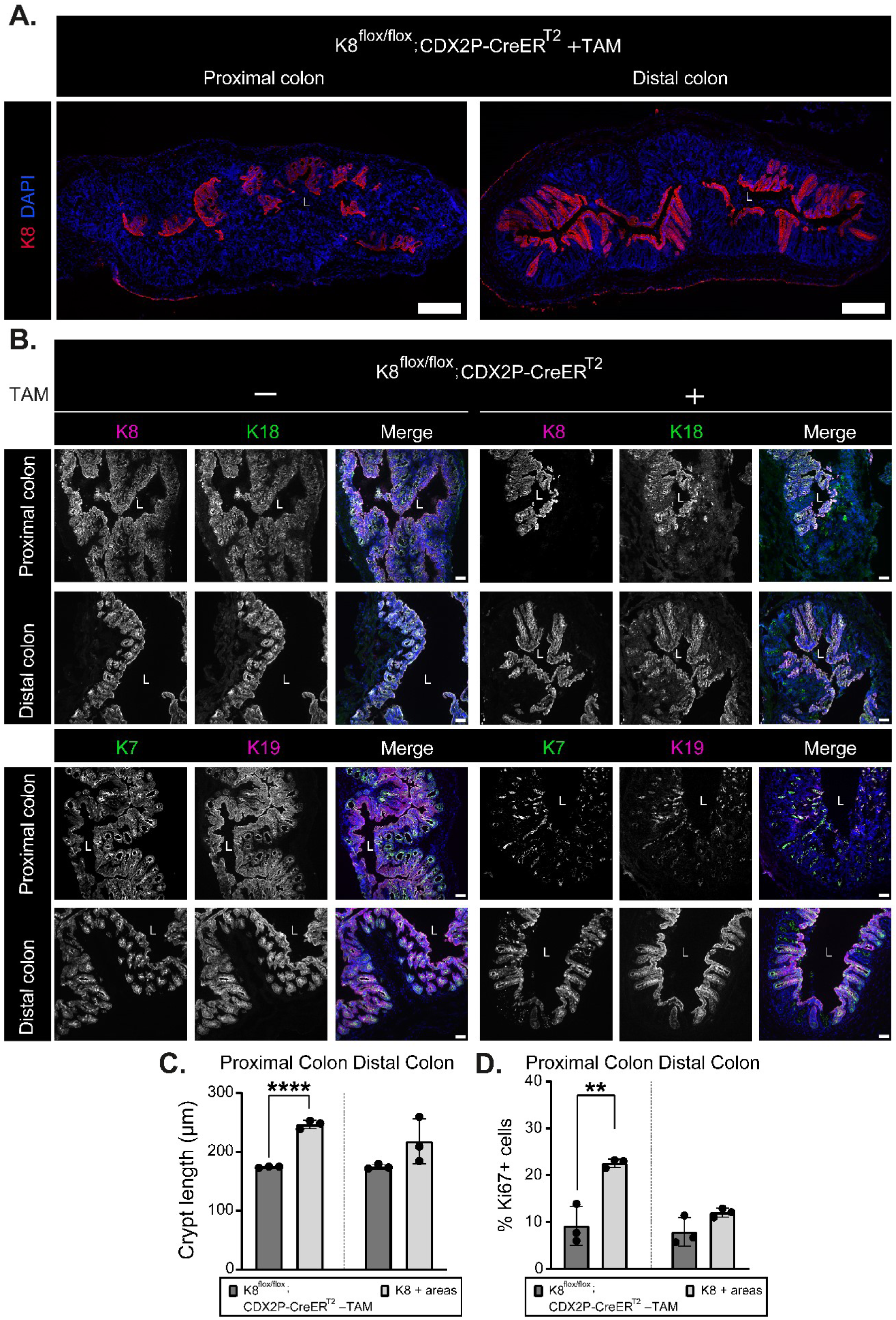
K8 downregulation in TAM-treated K8^flox/flox^; CDX2P-CreER^T2^ mice also induces patchy expression pattern of major other keratins in the colon. **A)** Representation of K8 (red) expression in entire proximal and distal colon section of K8^flox/flox^; CDX2P-CreER^T2^ +TAM mice (n=3 mice), nuclei, DAPI (blue) L=Lumen, Scale bar=200μm. **B)** Immunofluorescence staining of K8 (magenta) and K18 (green) or K19 (magenta) and K7 (green) with nuclei, DAPI (blue) in proximal and distal colon sections from K8^flox/flox^; CDX2P-CreER^T2^ (–TAM/+TAM) mice is shown, L=Lumen, Scale bar=50μm. **C)** Crypt lengths of K8^flox/flox^; CDX2P-CreER^T2^ –TAM and K8+ crypts of K8^flox/flox^; CDX2P-CreER^T2^ +TAM mice from proximal and distal colon were quantified and presented as mean (n=3 mice per group, 30 –TAM and 15 K8+ crypts per proximal/distal colon) ± SD, each data point represents an individual mouse. **D)** Percentage of Ki67+ cells in K8^flox/flox^; CDX2P-CreER^T2^ –TAM and K8+ crypts of K8^flox/flox^; CDX2P-CreER^T2^ +TAM mice from proximal and distal colon were quantified and presented as mean (n=3 mice per group, 3 images –TAM and 6–15 K8+ crypts per proximal/distal colon) ± SD, each data point represents an individual mouse. The statistical significance was determined after performing unpaired student’s t test for C and D, shown as * P < 0.05, ** P < 0.01, *** P < 0.001 and **** P < 0.0001.

**Supplemental Figure 2:**
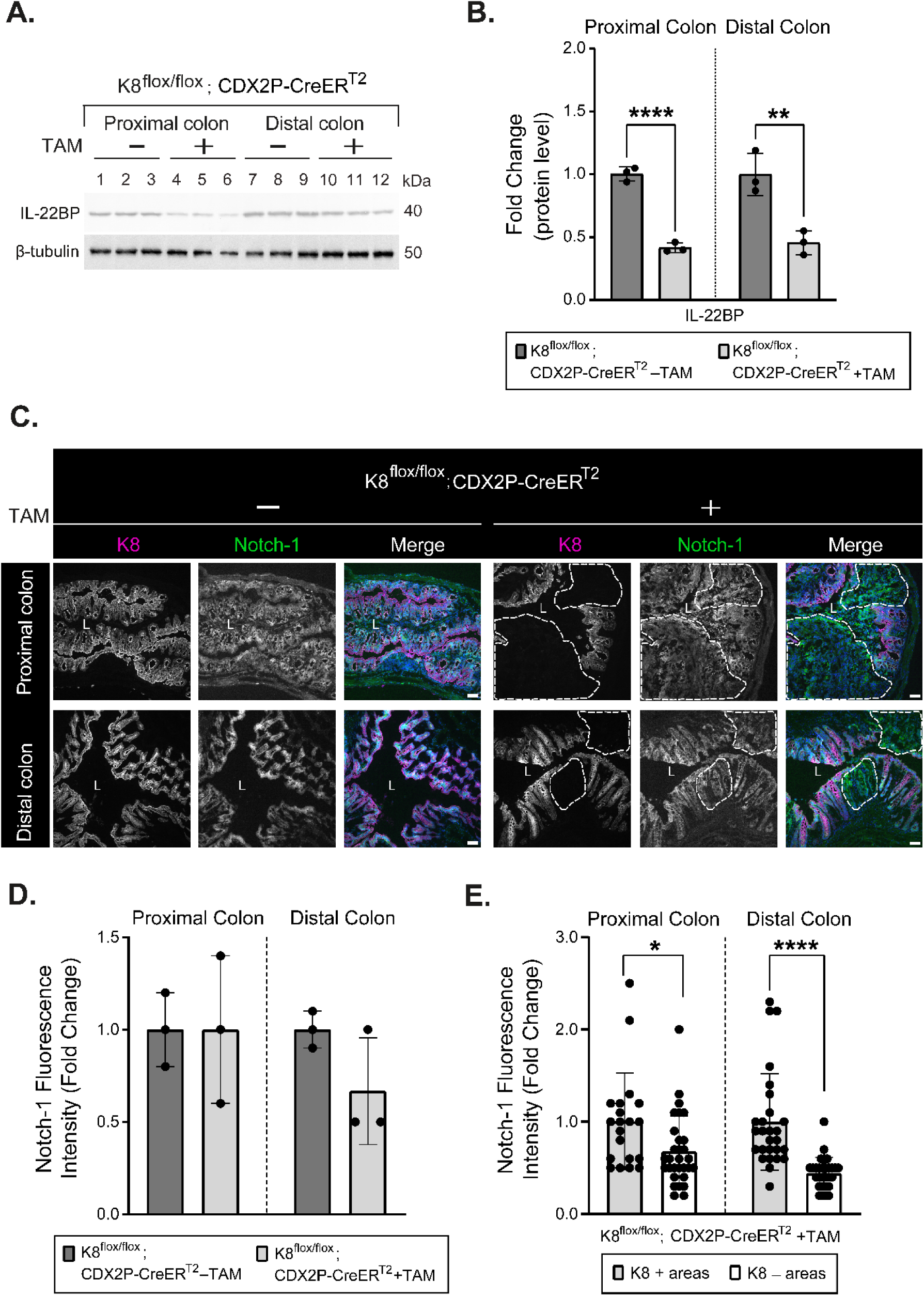
TAM-treated K8^flox/flox^; CDX2P-CreER^T2^ +TAM mice show decreased IL-22BP protein levels and reduced Notch-1 expression in the colon. **A)** Total proximal colon lysates from K8^flox/flox^; CDX2P-CreER^T2^ –TAM (Lanes 1–3), K8^flox/flox^; CDX2P-CreER^T2^ +TAM (Lanes 4–6) and total distal colon lysates from K8^flox/flox^; CDX2P-CreER^T2^ –TAM (Lanes 7–9), K8^flox/flox^; CDX2P-CreER^T2^ +TAM (Lanes 10–12) were immunoblotted for IL-22BP and β-tubulin was used as the loading control. **B)** Western blots from A were quantified and normalized to β-tubulin. The results are presented as the mean (n=3 mice per group, each data point represents an individual mouse) protein fold changes ± SD. **C)** Immunofluorescence staining of K8 (magenta), Notch-1 (green), nuclei, DAPI (blue) in proximal and distal colon sections of K8^flox/flox^; CDX2P-CreER^T2^ (–TAM/+TAM) mice (n=3 mice per group) is shown. Areas within the white dashed lines represent K8-negative colon crypts, L=Lumen, Scale bar=50μm. **D)** Mean fluorescence intensity for Notch-1 were quantified in proximal and distal colon of K8^flox/flox^; CDX2P-CreER^T2^ (–TAM/+TAM) mice and presented as mean fold change (n=3 mice per group, 2 images per proximal/distal colon) ± SD, each data point represents an individual mouse. **E)** Mean fluorescence intensity for Notch-1 in K8+ and K8– crypts of proximal and distal colon was measured in K8^flox/flox^; CDX2P-CreER^T2^ +TAM mice and presented as mean fold change (n=3 mice, 3–10 K8+ and 9–10 K8– crypts per proximal/distal colon) ± SD, each data point represents an individual crypt. The statistical significance was determined after performing unpaired student’s t test for B, D and E, shown as * P < 0.05, ** P < 0.01, *** P < 0.001 and **** P < 0.0001.

**Supplemental Figure 3:**
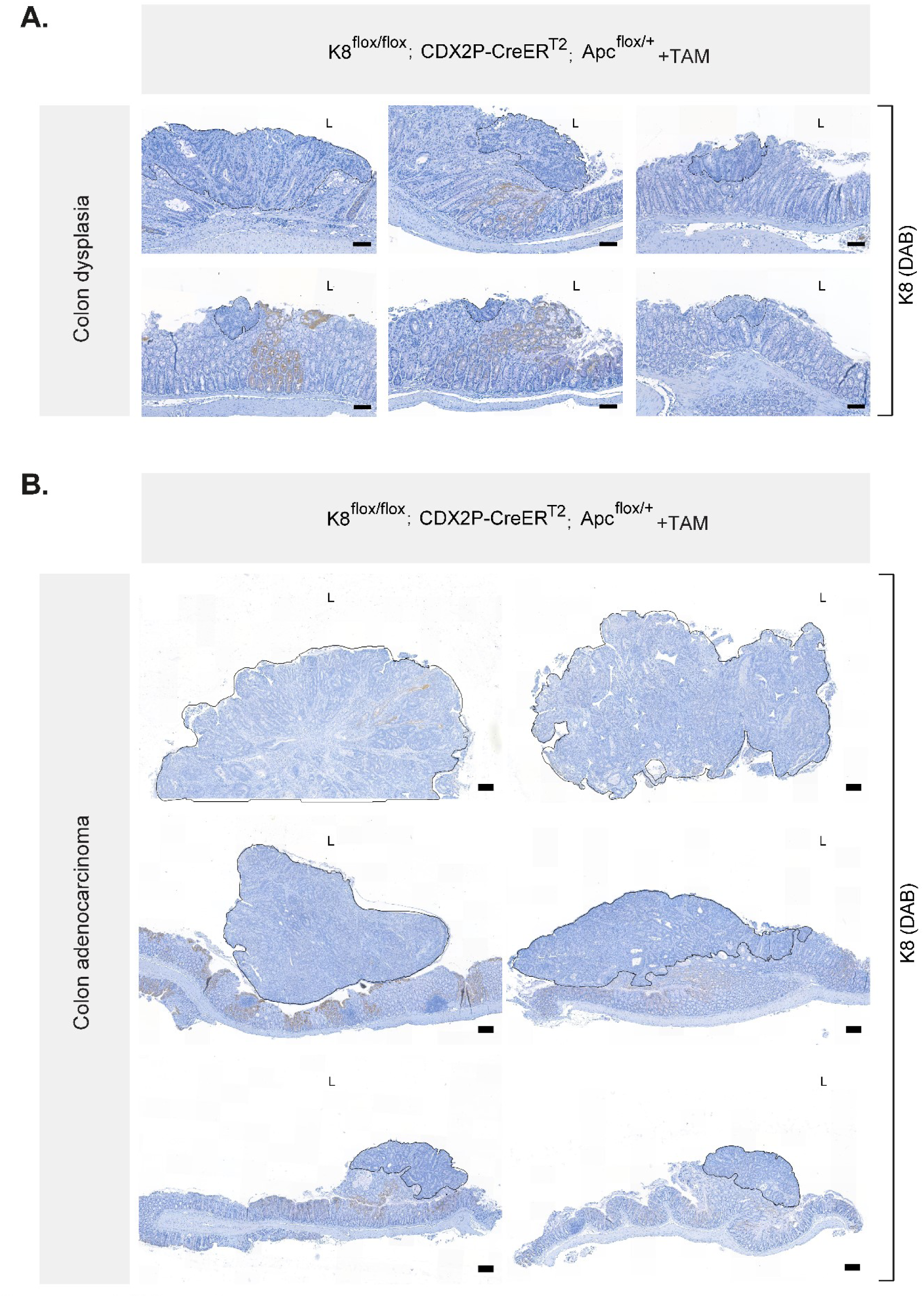
TAM-treated K8^flox/flox^; CDX2P-CreER^T2^; Apc^flox/+^ mice distal colon depict negligible K8 expression in the dysplastic areas and colon adenocarcinoma. **A–B)** K8 (DAB) immunolabeling in distal colon of K8^flox/flox^; CDX2P-CreER^T2^; Apc^flox/+^ mice, black encircled area in **A)** represent dysplastic growth regions in distal colon, L=Lumen, Scale bar=100 μm and in **B)** represent colon adenocarcinoma, L=Lumen, Scale bar=200 μm. < 1 % of all cells in these black encircled areas were K8-positive.

**Supplemental Figure 4:**
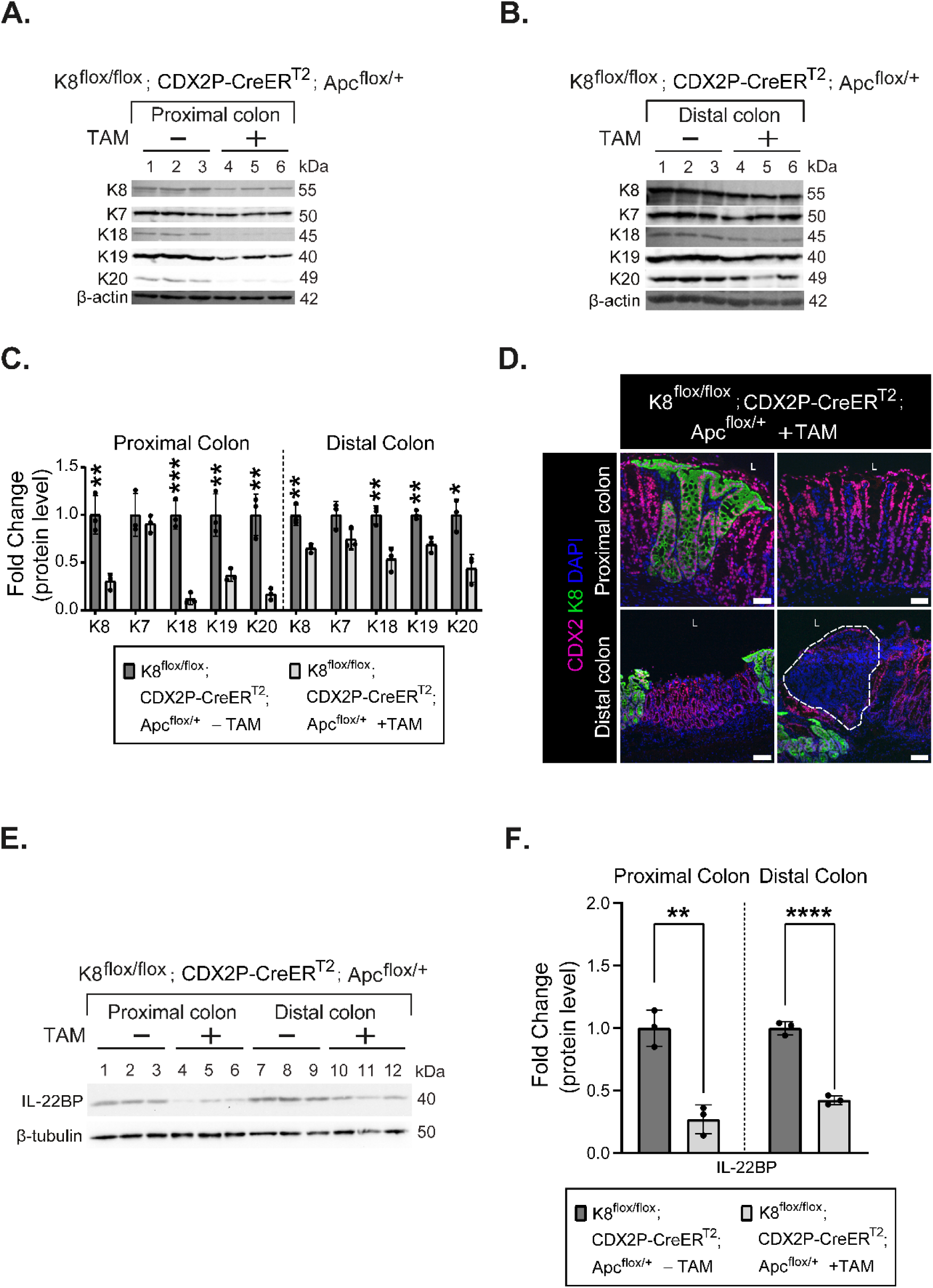
TAM-treated K8^flox/flox^; CDX2P-CreER^T2^; Apc^flox/+^ mice show downregulation of colonic keratins and reduced protein levels of IL-22BP in the colon. **A)** Total proximal colon lysates from K8^flox/flox^; CDX2P-CreER^T2^; Apc^flox/+^ –TAM (Lanes 1–3), K8^flox/flox^; CDX2P-CreER^T2^; Apc^flox/+^ +TAM (Lanes 4–6) and **B)** total distal colon lysates from K8^flox/flox^; CDX2P-CreER^T2^; Apc^flox/+^ –TAM (Lanes 1–3), K8^flox/flox^; CDX2P-CreER^T2^; Apc^flox/+^ +TAM (Lanes 4–6) were immunoblotted for K8, K7, K18, K19 and K20. β-actin was used as the loading control. **C)** Western blots from A and B were quantified and normalized to β-actin. The results are presented as mean (n=3 mice per group, each data point represents an individual mouse) protein fold changes ± SD. **D)** Immunofluorescence staining of K8 (green), CDX2 (magenta) and nuclei, DAPI (blue) in proximal and distal colon sections of K8^flox/flox^; CDX2P-CreER^T2^; Apc^flox/+^+TAM mice is shown, L=Lumen, Scale bar=50μm. Images are representative of n=2 mice. **E)** Total proximal colon lysates from K8^flox/flox^; CDX2P-CreER^T2^; Apc^flox/+^ –TAM (Lanes 1–3), K8^flox/flox^; CDX2P-CreER^T2^; Apc^flox/+^ +TAM (Lanes 4–6) and total distal colon lysates from K8^flox/flox^; CDX2P-CreER^T2^; Apc^flox/+^ –TAM (Lanes 7–9), K8^flox/flox^; CDX2P-CreER^T2^; Apc^flox/+^ +TAM (Lanes 10–12) were immunoblotted for IL-22BP and β-tubulin was used as the loading control. **F)** Western blots from E were quantified and normalized to β-tubulin. The results are presented as the mean (n=3 mice per group, each data point represents an individual mouse) protein fold changes ± SD. The statistical significance was determined after performing unpaired student’s t test for C and F, shown as * P < 0.05, ** P < 0.01, *** P < 0.001 and **** P < 0.0001.

